# The genome sequence of the hemlock woolly adelgid, *Adelges tsugae* Annand 1924

**DOI:** 10.1101/2025.04.07.644581

**Authors:** Bryan M.T. Brunet, Dustin T. Dial, Gaelen R. Burke, Carol D. von Dohlen, Julia Douglas Freitas, Haley Sanderson, Afiya R. Chida, Samantha J. Jones, Fergal J. Martin, Leanne Haggerty, Stephen W. Scherer, Ioannis Ragoussis, Steven J.M. Jones, Robert G. Foottit, Nathan P. Havill

## Abstract

The genome of the hemlock woolly adelgid (*Adelges tsugae*) is presented in 10 chromosomal pseudomolecules along with 132 unplaced scaffolds for a combined length of 216.56 Mb. The genome is highly contiguous, with a benchmarking universal single-copy orthologs (BUSCO) completion score of 98.25%, an N50 of 20.54 Mb, and the longest scaffold 37.41 Mb in length. On the basis of chromosomal synteny with other aphid genomes, the hemlock woolly adelgid’s genome is composed of 8 autosomes and two sex (X) chromosomes. Annotation of the assembly identified 11,800 coding genes, 1,930 non-coding genes, and 20,403 mRNA transcripts. The mitogenome is also presented in a single, annotated, circular contig 25,980 bases in length.

**Species taxonomy:** Eukaryota; Metazoa; Arthropoda; Hexapoda; Insecta; Pterygota; Neoptera; Paraneoptera; Hemiptera; Sternorrhyncha; Aphidomorpha; Adelgoidea; Adelgidae; *Adelges*; *Annandina*; *Adelges* (*Annandina*) *tsugae* Annand, 1924 (NCBI:txid357502)

## Introduction

Few adventive aphids (Hemiptera: Sternorrhyncha: Aphidomorpha) are as economically important to North American forests as the hemlock woolly adelgid (HWA), *Adelges tsugae* Annand 1924. This species has devastating and disruptive effects on native eastern hemlock species (*Tsuga canadensis* (Linnaeus) Carrière and *T. carolinensis* Engelmann) – foundation species that provide vital ecosystem services in eastern forests^1^ – and their biotic and abiotic associates, including the alteration of forest vegetation composition, declines in associated fauna, and ecosystem-level changes^2^. HWA was introduced to North America in or near Richmond, Virginia before 1951 from its native range in southern Japan^3^ and has since spread at a rate of 2.5-37 km per year to cover most of the native hemlock forest in the eastern United States^4, 5, 6, 7^. It is now also threatening hemlocks in eastern Canada, with recent detections in southern Ontario^8, 9^ and an established population in Nova Scotia^10, 11^. Moreover, species distribution models forecast that HWA will infest all but a narrow band of hemlock forest north of the 45th parallel by 2050^2, 12^.

Unlike other significant insect forest pests in North America^13, 14^ (*e.g.,* Mountain Pine Beetle^15^, Asian Longhorned Beetle^16^, Spotted Lanternfly^17^, Emerald Ash Borer^18^, Spongy Moth^19, 20^, and White Pine Weevil^21^), robust genomic resources remain to be developed for HWA. In fact, among the Aphidomorpha (including the Adelgidae, Phylloxeridae, and Aphididae), only the Aphididae and Phylloxeridae are represented among species with available chromosome-length genome assemblies, and at present, the only adelgid with a scaffold-level assembly is the Cooley spruce gall adelgid, *Adelges cooleyi* (Gillette, 1907)^22^. High-quality genome assemblies provide a foundation for understanding the genetic architecture of complex traits, and are an important resource in the molecular arsenal used in the development of management and biocontrol strategies^16, 18, 19, 23, 24, 25, 26,27^. Here, we present a chromosome-scale genome assembly for HWA, the first of this magnitude for the Adelgidae, to bolster resources available for future work on the comparative genomics of the Aphidomorpha and the genetic basis of life history traits that allow HWA to exploit new areas in its invasive range.

## Methods

### Sample collection and extraction

Whole adults, nymphs, and/or eggs of HWA were collected by hand on five occasions between 2013 and 2021. Samples were reared for up to one week at room temperature to accumulate specimens before transferring to 95% ethanol and storing at -80°C until further use. Four samples were taken from previously identified HWA-infested eastern hemlock (*T. canadensis*) at the USDA Forest Service Northern Research Station in Hamden, New Haven County, Connecticut, and one from another location 4 km away.

Collection information associated with each collection event is presented in **Table 1**. A slide-mounted adult voucher specimen of sample ihAdeTsug4 was retained and deposited in the Yale Peabody Museum (YPM; Connecticut, USA) with accession number YPM#ENT595371. Previously unpublished Illumina short-read sequence data for sample ihAdeTsug1 was also used. This sample was the source for the two previously published endosymbiont genomes^28^.

**Table 1:** Sample collection data with assigned Tree of Life ID (ToLID) and Canada Biogenome Project sample number.

| ToLID | Specimen ID (CBP#) | Collection Locality | Date | Collector |
| --- | --- | --- | --- | --- |
| ihAdeTsug1 | 13-050 (CBP02084) | USA: Connecticut, New Haven County, Hamden, USDA Forest Service Northern Research Station, *Tsuga canadensis* (41.3847, -72.917) | 2013-04-22 | N.P. Havill |
| ihAdeTsug2 | 1202-1 (CBP02085) | USA: Connecticut, New Haven County, Hamden, USDA Forest Service Northern Research Station, *Tsuga canadensis* (41.3847, -72.917) | 2017-11-29 | N.P. Havill |
| ihAdeTsug3 | 18-687-01 (CBP02086) | USA: Connecticut, New Haven County, Hamden, USDA Forest Service Northern Research Station, *Tsuga canadensis* (41.3847, -72.917) | 2018-12-16 | N.P. Havill |
| ihAdeTsug4 | 19-124-03 (CBP02087) | USA: Connecticut, New Haven County, Hamden, Deerfield Rd., *Tsuga canadensis* (41.4246, -72.9329) | 2019-06-12 | M.E. Montgomery |
| ihAdeTsug5 | 21-098 (CBP02088) | USA: Connecticut, New Haven County, Hamden, USDA Forest Service Northern Research Station, *Tsuga canadensis* (41.3847, -72.917) | 2021-01-13 | N.P. Havill |

Isolation of DNA from each sample was performed on pooled specimens using commercially available kits, including the Qiagen Blood and Cell Culture DNA Mini kit (ihAdeTsug2) and MagAttract HMW DNA kit (ihAdeTsug3 and ihAdeTsug4). Since eastern North American populations of HWA reproduce parthenogenetically and likely represent a single invasive clone throughout its introduced range^3^, pooling of specimens within samples, as well as the use of multiple distinct samples collected at different times (**Table 1**), does not pose a challenge to genome assembly as might be expected with sampling individuals of differing genetic backgrounds.

### Library construction and sequencing

DNA from two samples, ihAdeTsug3 and ihAdeTsug4, were selected for long-read sequencing. Fragment length analysis identified peaks in the fragment length distributions for these extractions at 16 Kb and 11 Kb, respectively, so a Pacific Biosciences long-insert library was prepared by combining the samples and selecting for fragments of 15 Kb in size without shearing at the University of Maryland School of Medicine’s Institute for Genome Sciences, Genome Resource Centre (Maryland, USA). Continuous long-read sequencing was performed in a single SMRT Cell 8M run on a PacBio Sequel II over a 15-hour movie collection time. A 10x Genomics Chromium linked-read library was also prepared using sample ihAdeTsug2, and the sample was subsequently paired-end (2 x 150 bp) sequenced on an Illumina HiSeq X at the McGill University and Génome Québec Innovation Centre (Québec, Canada). Finally, sample ihAdeTsug5 was used to prepare a single Arima Hi-C sequencing library, which was then paired-end (2 x 150 bp) sequenced in two runs on an Illumina MiSeq v2 at the McGill Genome Centre (Québec, Canada). The Illumina (2 x 150 bp with 500 bp inserts) libraries for sample ihAdeTsug1 noted above were previously prepared and sequenced on a HiSeq 2500 at the Yale Centre for Genome Analysis^28^ (Connecticut, USA).

Genome size was determined through flow cytometric estimation using genomic DNA isolated from two adult females following standard protocols for insects^29^. *Drosophila virilis* (1C = 328 Mb) was used as the standard.

### Genome assembly

Raw PacBio subreads were input to Falcon/Falcon-Unzip^30^ (v.1.3.7) using the PacBio Assembly Tool Suite (pb-assembly v.0.0.8) installed via the Conda package and environment management system. Parameters used in the assembly and subsequent steps are defined in **Table 2**. The full Longstitch pipeline^31^ (v.1.0.5) was used to correct mis-assembled contigs and then scaffold and gap-fill using pre-assembly reads (preads) from the Falcon output. Default parameters were used, other than the minimum number of spanning molecules required for a correct assembly and the maximum distance allowed between reads aligned within the same molecule, which were determined automatically by defining the genome size. Three rounds of gap-filling and scaffolding were also performed at the ntLink^32^ (v.1.3.9) step.

**Table 2:**
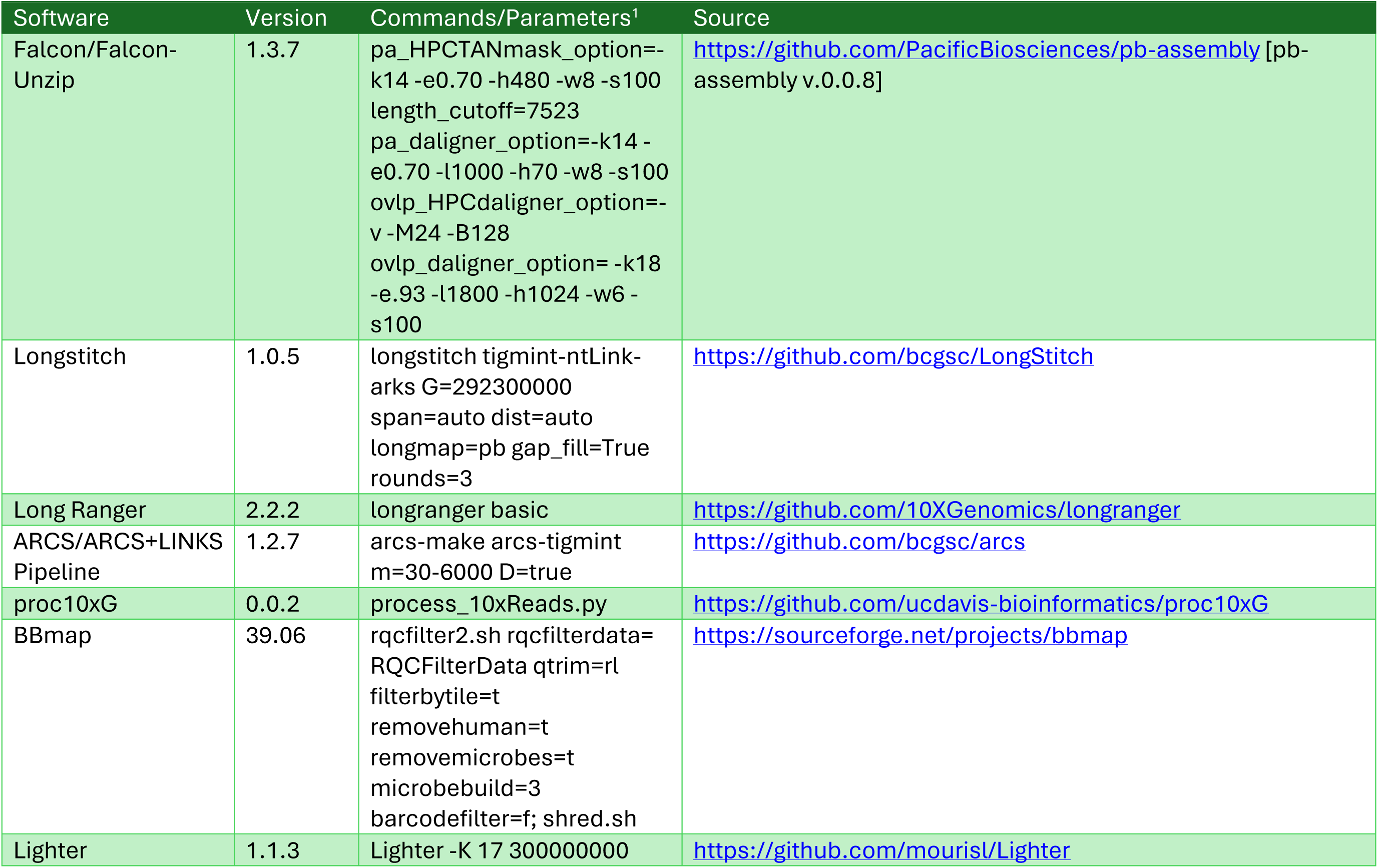
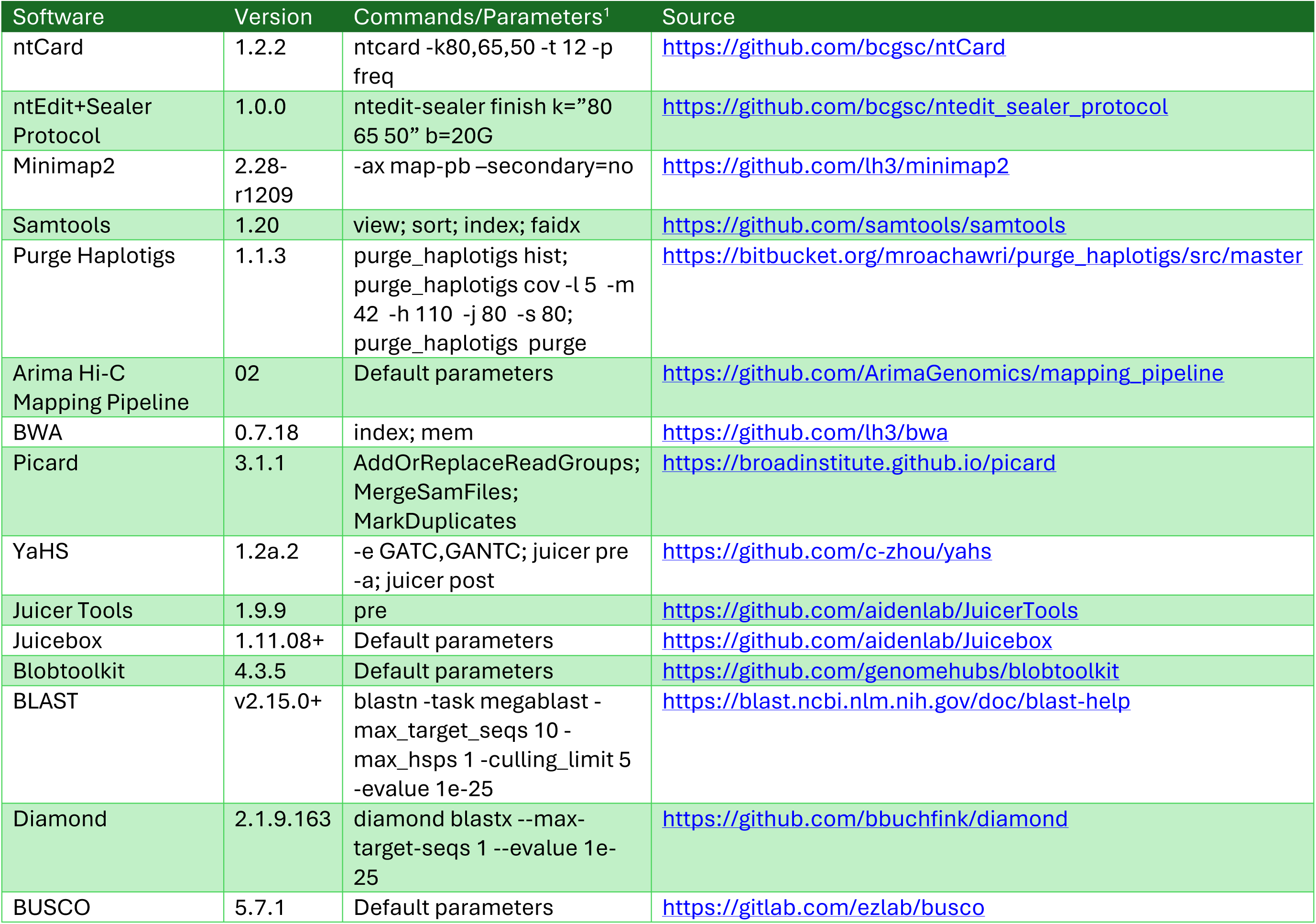
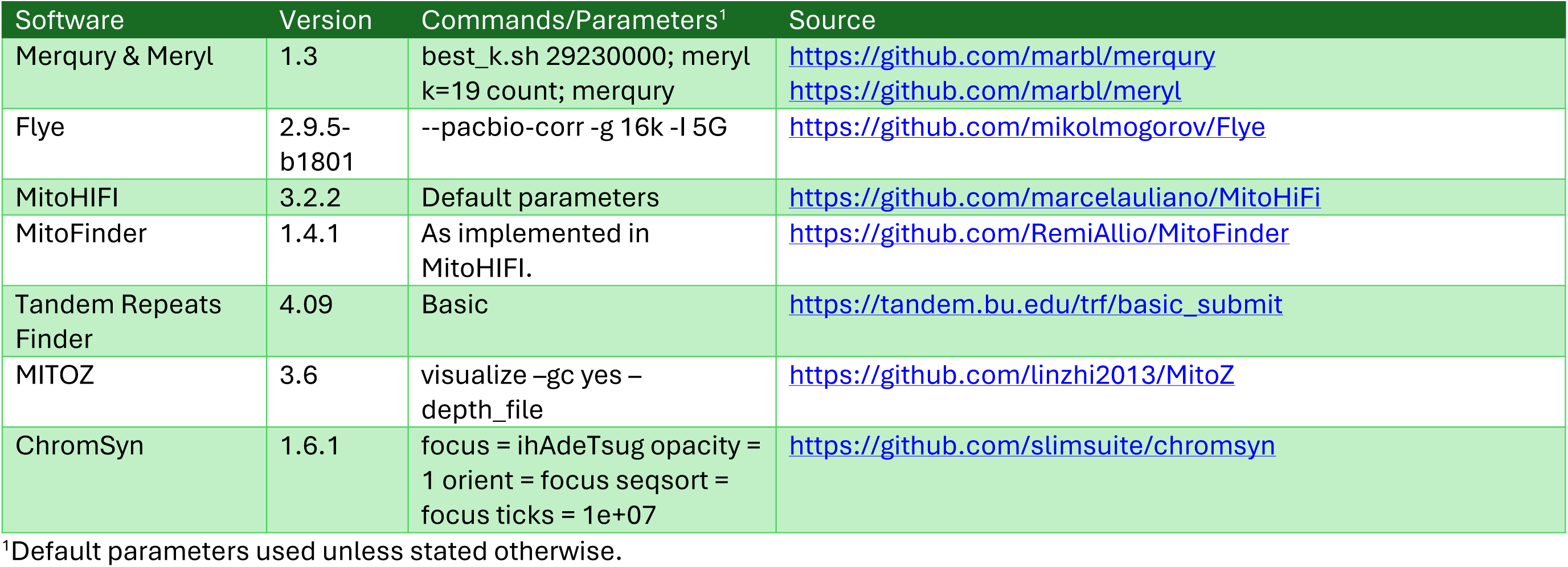
Software tools used by order of processing.

The ARCS+LINKS pipeline, as implemented in ARCS^33^ (v.1.2.7) with Tigmint^34^ (v.1.2.10) and LINKS^35^ (v.2.0.1), was used to identify and correct misassembled scaffolds using 10x Chromium linked-reads. The pipeline was run using the ‘arcs-tigmint’ command with distance estimation enabled and the linked-read barcode multiplicity range set to 30-6000 to ignore both low and high outliers in the multiplicity distribution. Linked-reads were first barcode-processed using Long Ranger’s^36^ (v.2.2.2) ‘basic’ command as required for input into ARCS.

Short-read polishing and gap-filling was completed using the ntEdit+Sealer Assembly Finishing Protocol^37^ (v.1.0.0). Illumina short-reads and 10x linked-reads were trimmed and filtered of low-quality and contaminant sequences using RQCFilter2 in BBmap^38^ (v.39.06), and then error-corrected using Lighter^39^ (v.1.1.3) before being input into ntEdit+Sealer. Lighter was run with automated estimation of alpha using a *k*-mer length of 17 and an approximated genome size of 300 Mb. Prior to trimming and filtering, the 10x linked-reads were first barcode-processed using process_10xReads.py (v.0.0.2) of proc10xG (https://github.com/ucdavis-bioinformatics/proc10xG). The Bloom filter size used in ntEdit+Sealer was estimated at 20 GB using ntCard^40^ (v.1.2.2).

Redundant and junk scaffolds were then identified and removed from the assembly using Purge Haplotigs^41^ (v.1.1.3) following alignment of the raw PacBio subreads to the assembly. Low-, mid-, and high-read depth cutoffs were estimated at 5, 42, and 110 from the coverage histogram produced using the ‘hist’ command.

Arima Hi-C reads were mapped to the purged assembly, and filtered on the basis of chimerism and mapping quality using the Arima mapping pipeline (v.02, https://github.com/ArimaGenomics/mapping_pipeline). Technical replicates were also merged before removal of duplicate reads. Hi-C scaffolding was performed with YaHS^42^ (v.1.2a.2) using the Arima genomics two-enzyme combination. Juicer Tools^43^ (v.1.9.9) was used to generate a Hi-C contact matrix for visualization in Juicebox^44^ (v.1.11.08+). Manual curation of the assembly was performed using Juicebox Assembly Tools^45^.

The curated assembly was checked for contamination using Blobtools2 from the Blobtoolkit^46^ (v.4.3.5). First, each scaffold was partitioned into 100 Kb sequences with 500 bp overlap using the shred.sh script from BBmap, and then each partition was queried against the NCBI nt database (released 05-July-2024) using blastn^47^ (v2.15.0+). Similarly, Diamond blastx^48^ (v.2.1.9.163) searches were performed against the UniProt^49^ reference proteomes (release 2024_06) database. Coverage of the raw PacBio subreads was then assessed by mapping them to the genome assembly using minimap2^50^ (v.2.28-r1209). BUSCO^51^ (v.5.7.1) was used to assess the completeness of the assembly using the Hemiptera_odb10 dataset. Finally, the fraction of *k*-mers that are truly represented in the genome and not resulting from sequencing error (*k*-mer completeness) as well as the frequency of sequence consensus errors (assembly consensus quality, QV) were calculated with Merqury^52^ (v.1.3) using the adapter-trimmed, contaminant-filtered, and error-corrected 10x linked-reads after preparation of a meryl *k*-mer database (k=19).

Since the mitogenome was not identified among the final set of scaffolds, it was assembled separately. Pre-assembly reads from the Falcon assembly noted above were mapped to a publicly available mitogenome reference of HWA collected in Taiwan^53^ (MT263947) using minimap2. Mapped reads were assembled into contigs using Flye^54^ (v.2.9.5-b1801), and then contigs were used as input for MitoHiFi^55^ (v.3.2.1), where they were subsequently circularized and annotated using MitoFinder^56^ (v.1.4.1). Tandem Repeats Finder^57^ (v.4.09) was used to identify tandem repeats signaling the putative control and repeat regions in non-coding locations of the mitogenome. The mitogenome was then visualized using MITOZ^58^ (v.3.6).

### Chromosomal synteny

ChromSyn^59^ (v.1.6.1) was used to identify sex chromosomes and chromosomal rearrangements between HWA and other aphid species with chromosome-scale assemblies using syntenic regions sharing complete single-copy orthologs from the BUSCO Hemiptera_odb10 dataset. Namely, comparisons were made with two species in which the sex chromosomes were previously identified, including horned-gall aphid *Schlechtendalia chinensis* (Bell, 1851) (Aphididae: Eriosomatinae) ^60^ (GCA_019022885.1) and corn leaf aphid, *Rhopalosiphum maidis* (Fitch, 1856) (Aphididae: Aphidinae) ^61^ (GCF_003676215.2).

## Results

### Genome sequence report

The genome of the hemlock woolly adelgid was assembled from multiple samples collected in Hamden Connecticut, USA, together representing the single multi-locus lineage that predominates the northeast US and Canada^3^. The genome was sequenced to a total coverage of 101-fold using PacBio continuous long-reads based on an estimate of 292.3 ± 1.1 Mb (n=2) for the genome size using flow cytometry. In addition to the long-read data, newly generated 10x linked-reads and Hi-C data provided 249-fold and 24-fold raw coverage, respectively, and the short-read data^28^ provided an additional 86-fold coverage. The resulting assembly consisted of 145 scaffolds. Following manual curation, the assembly was reordered to accommodate two sex chromosomes (X1 and X2) identified through comparisons of chromosomal synteny with *R. maidis* and *S. chinensis*, as well as to merge chromosome X1 with two smaller unplaced scaffolds through translocation corrections (**Figure 1**). A 5 Kb gap associated with one of these corrections was also excised and subsequently removed from the assembly, along with one other scaffold that was less than 200 bp in length, leaving the final scaffold count at 142.

**Figure 1.**
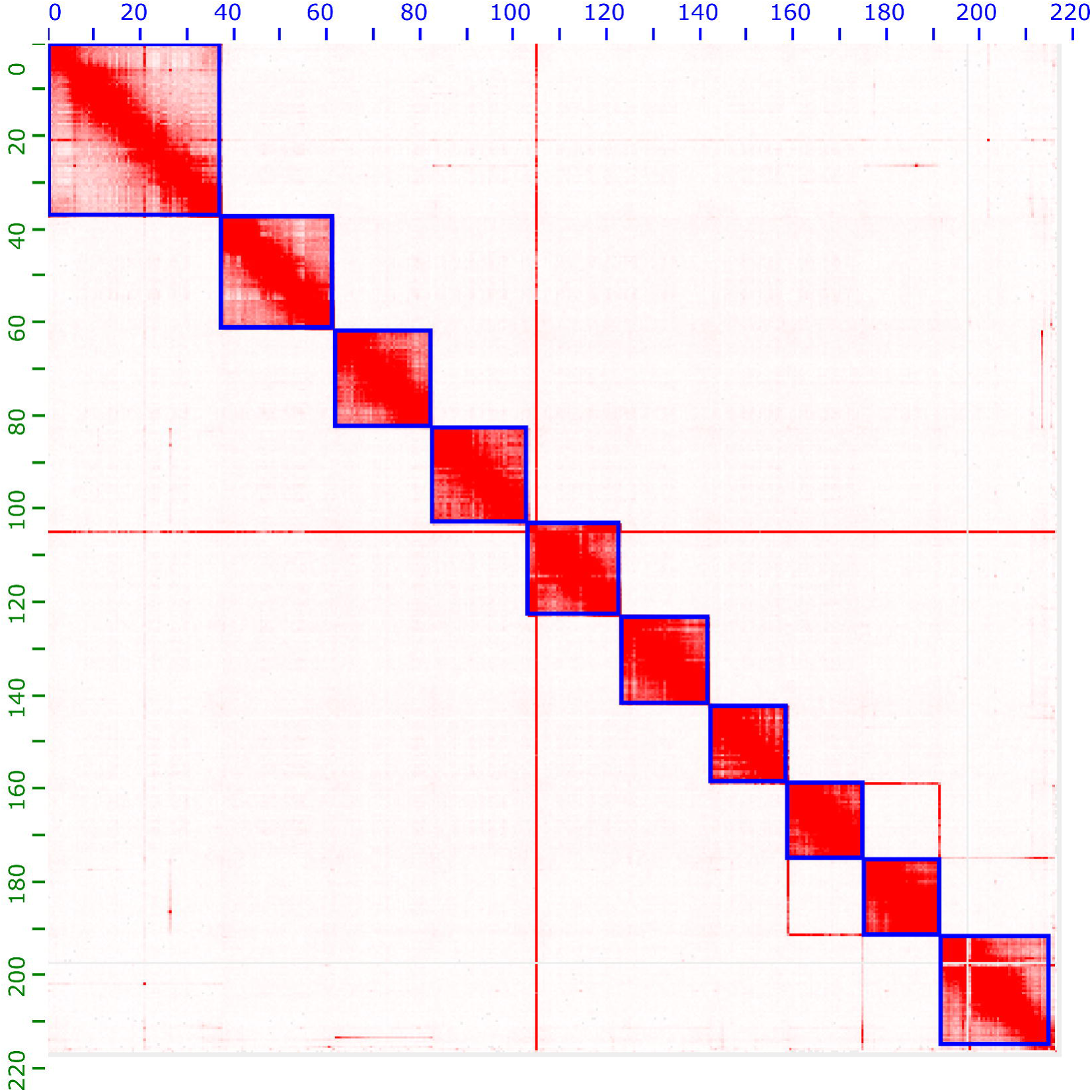
Genome assembly of *Adelges tsugae*, ihAdeTsug: Hi-C contact map. Hi-C contact map of *A. tsugae* (ihAdeTsug) assembly visualized using JuiceBox with Assembly Tools. Autosomes are shown in order of size from left to right and top to bottom, with chromosomes X2 and then X1 following autosome 8. Unplaced scaffolds 1-132 are visible at the lower right.

The final curated assembly has a total length of 216.56 Mb, with the longest scaffold at 37.41 Mb and an N50 of 20.54 Mb (**Table 3**, **Figure 2**). In addition, over 99% of the HWA genome was recovered in just 10 chromosome-scale pseudomolecules – each greater than 16 Mb in length – representing the expected haploid number of chromosomes for this species based on comparisons to other adelgids^62^ (**Table 4**, **Figure 3**). All scaffolds with significant hits to homologues in the NCBI nt and Uniprot reference proteomes databases, matched to representatives from insect families, with the Adelgidae (215.19 Mb) and Aphididae (835.50 Kb) accounting for 58 scaffolds and a combined length of 216.03 Mb (**Figure 4**). The *k*-mer completeness and QV were 87.2% and 44.1, respectively, indicating a highly complete and error-free assembly. The Merqury spectra-cn plot (**Figure 5**) shows low *k*-mer duplication, and a large diploid peak at a multiplicity of 140 with a nearly undetectable haploid peak at 75. This suggests strong homozygosity, as would be expected for the invasive eastern North American *A. tsugae* lineage which reproduces only by parthenogenesis and experienced a strong genetic bottleneck when introduced^3^.

**Table 3:**
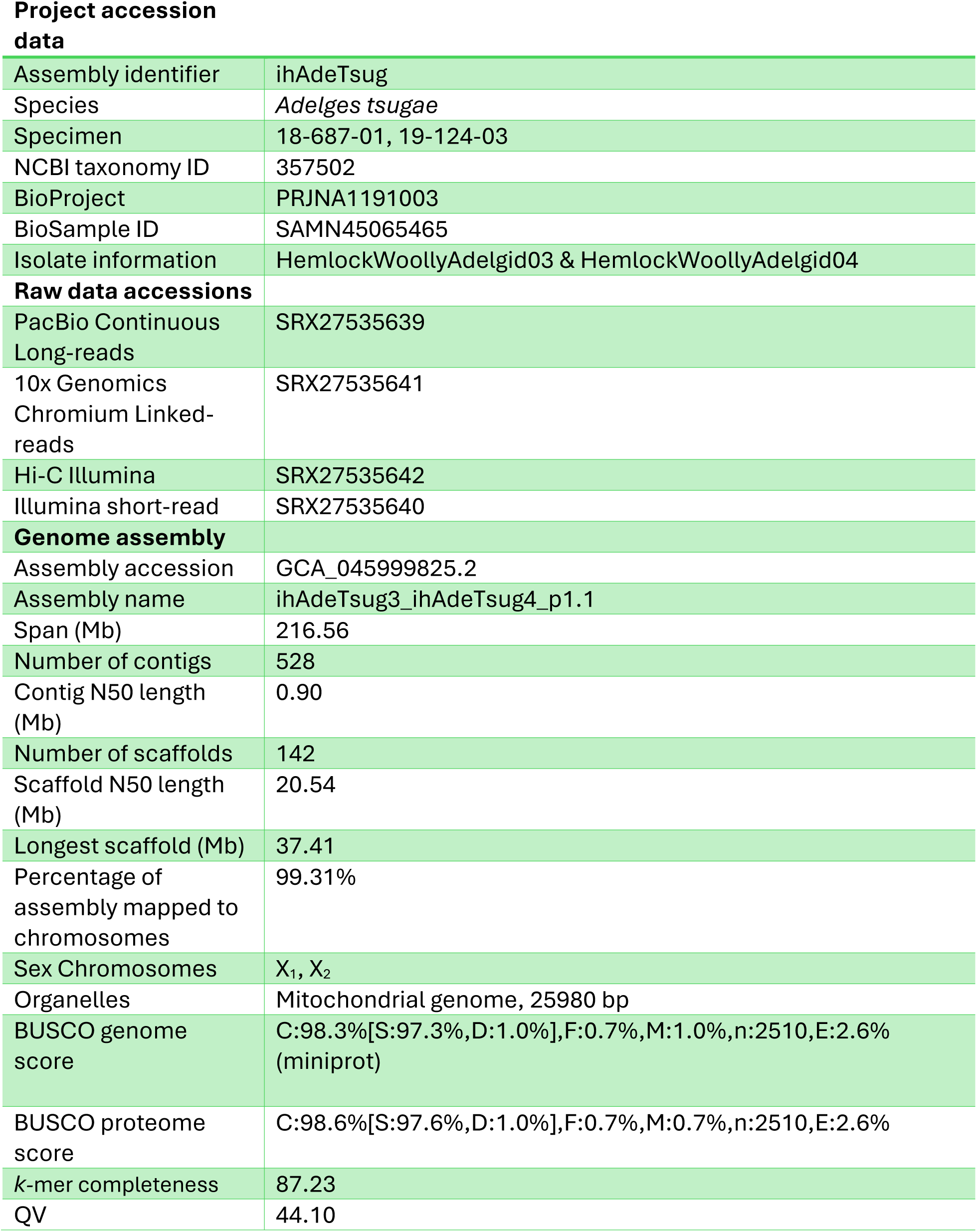
Genome data for *Adelges tsugae*, ihAdeTsug.

|  |  |
| --- | --- |
| **Project accession data** |  |
| Assembly identifier | ihAdeTsug |
| Species | *Adelges tsugae* |
| Specimen | 18-687-01, 19-124-03 |
| NCBI taxonomy ID | 357502 |
| BioProject | PRJNA1191003 |
| BioSample ID | SAMN45065465 |
| Isolate information | HemlockWoollyAdelgid03 & HemlockWoollyAdelgid04 |
| **Raw data accessions** |  |
| PacBio Continuous Long-reads | SRX27535639 |
| 10x Genomics Chromium Linked-reads | SRX27535641 |
| Hi-C Illumina | SRX27535642 |
| Illumina short-read | SRX27535640 |
| **Genome assembly** |  |
| Assembly accession | GCA_045999825.2 |
| Assembly name | ihAdeTsug3_ihAdeTsug4_p1.1 |
| Span (Mb) | 216.56 |
| Number of contigs | 528 |
| Contig N50 length (Mb) | 0.90 |
| Number of scaffolds | 142 |
| Scaffold N50 length (Mb) | 20.54 |
| Longest scaffold (Mb) | 37.41 |
| Percentage of assembly mapped to chromosomes | 99.31% |
| Sex Chromosomes | X<sub>1</sub>, X<sub>2</sub> |
| Organelles | Mitochondrial genome, 25980 bp |
| BUSCO genome score | C:98.3%[S:97.3%,D:1.0%],F:0.7%,M:1.0%,n:2510,E:2.6% (miniprot) |
| BUSCO proteome score | C:98.6%[S:97.6%,D:1.0%],F:0.7%,M:0.7%,n:2510,E:2.6% |
| k-mer completeness | 87.23 |
| QV | 44.10 |

**Figure 2.**
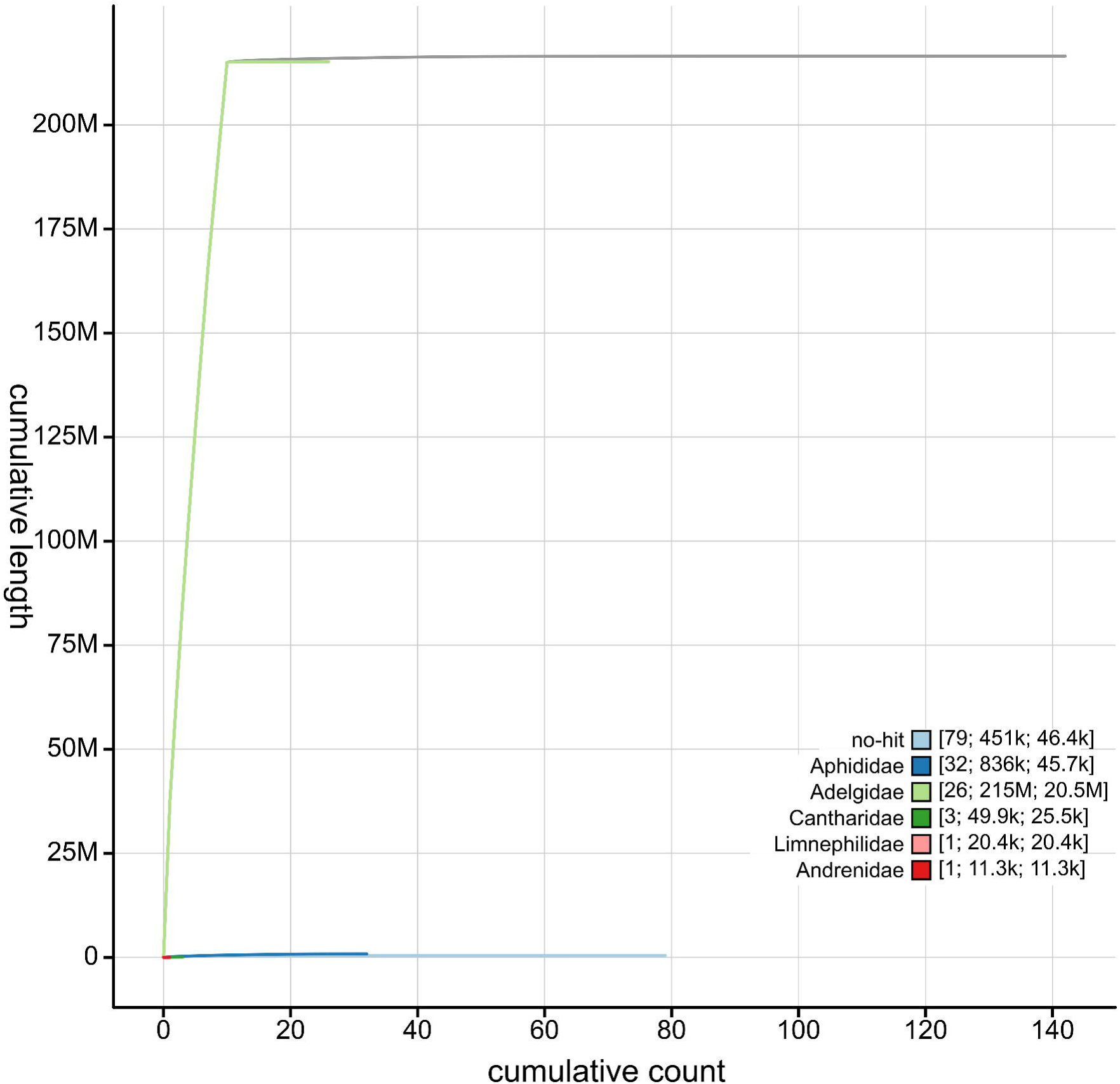
Genome assembly of *Adelges tsugae*, ihAdeTsug: cumulative sequence length. BlobToolKit cumulative sequence plot of *A. tsugae* (ihAdeTsug) assembly. The grey line shows cumulative length for all scaffolds. Coloured lines show cumulative lengths of scaffolds assigned to each family using the bestsumorder taxrule.

**Table 4:** Chromosomal pseudomolecules in the genome assembly of *Adelges tsugae,* ihAdeTsug.

| GenBank accession | Chromosome | Size (Mb) | GC content % |
| --- | --- | --- | --- |
| CM099565.1 | 1 | 37.41 | 30.5 |
| CM099566.1 | 2 | 24.27 | 30.5 |
| CM099567.1 | 3 | 21.14 | 30.5 |
| CM099568.1 | 4 | 20.53 | 31 |
| CM099569.1 | 5 | 19.74 | 31.5 |
| CM099570.1 | 6 | 19.20 | 30.5 |
| CM099571.1 | 7 | 16.66 | 31.5 |
| CM099574.1 | 8 | 16.10 | 31.5 |
| CM099573.1 | X1 | 23.46 | 31 |
| CM099572.1 | X2 | 16.55 | 31 |
| CM099575.1 | MT | 25,980<sup>1</sup> | 14.5 |

<sup>1</sup>Mitogenome size is listed in bases.

**Figure 3.**
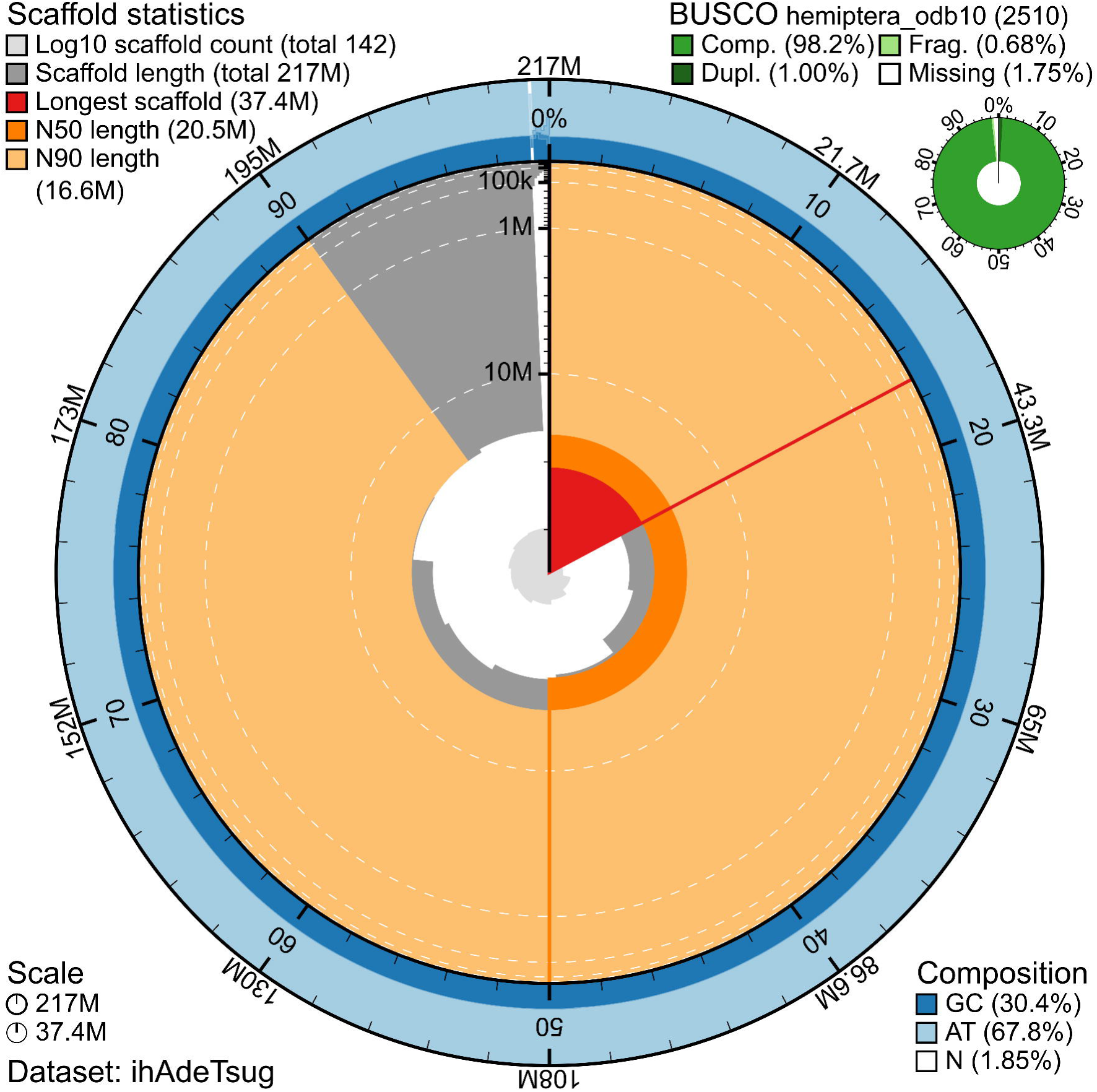
Genome assembly of *Adelges tsugae*, ihAdeTsug: metrics. BlobToolKit snail plot showing N50 metrics, base pair composition, and BUSCO gene completeness for *A. tsugae* (ihAdeTsug) assembly. The main plot is divided into 1,000 size-ordered bins around the circumference, with each bin representing 0.1% of the 216,556,762 bp assembly. The distribution of chromosome lengths is shown in dark grey, and the plot radius scaled to the longest chromosome present in the assembly (37,413,131 bp) is shown in red. Orange and pale-orange arcs show the N50 and N90 chromosome lengths (20,533,966 and 16,550,355 bp, respectively). The pale grey spiral shows the cumulative chromosome count on a log scale with white scale lines showing successive orders of magnitude. The blue and pale-blue area around the outside of the plot displays the distribution of GC (blue), AT (pale blue) and N (white) percentages, using the same bins as the inner plot. A summary of complete (98.2%), fragmented (0.68%), duplicated (1.0%), and missing (1.75%) BUSCO genes in the hemiptera_odb10 set is shown in the top right.

**Figure 4.**
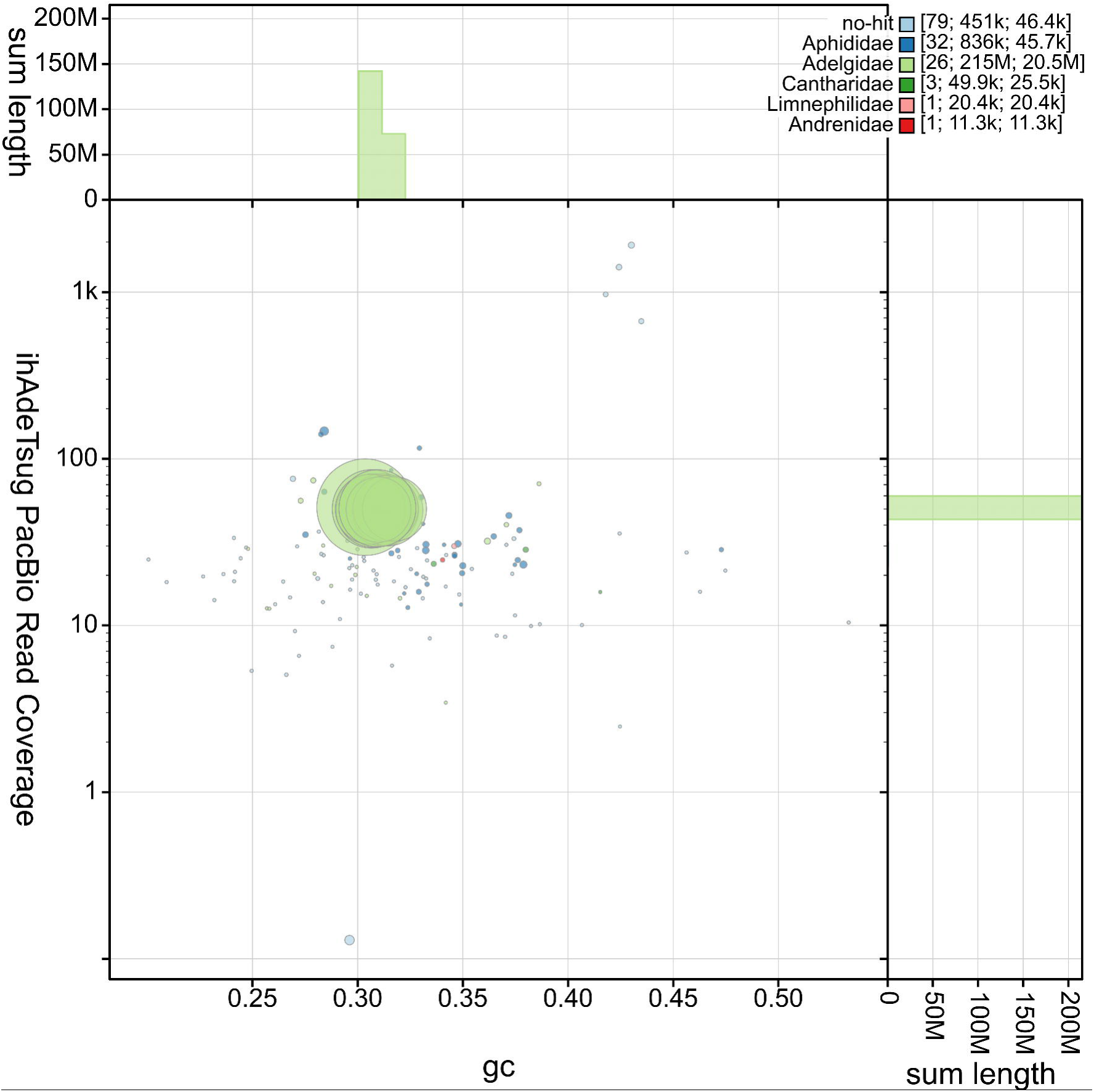
Genome assembly of *Adelges tsugae*, ihAdeTsug: GC coverage. BlobToolKit GC coverage plot of *A. tsugae* (ihAdeTsug) assembly. Scaffolds are coloured by family with Adelgidae represented by light green, Aphididae by blue, and no-hits represented by pale blue. Circles are sized in proportion to scaffold length. Histograms show the distribution of scaffold length sum along each axis. Legend at the top right shows the number of matching scaffolds, cumulative sequence length, and N50.

**Figure 5.**
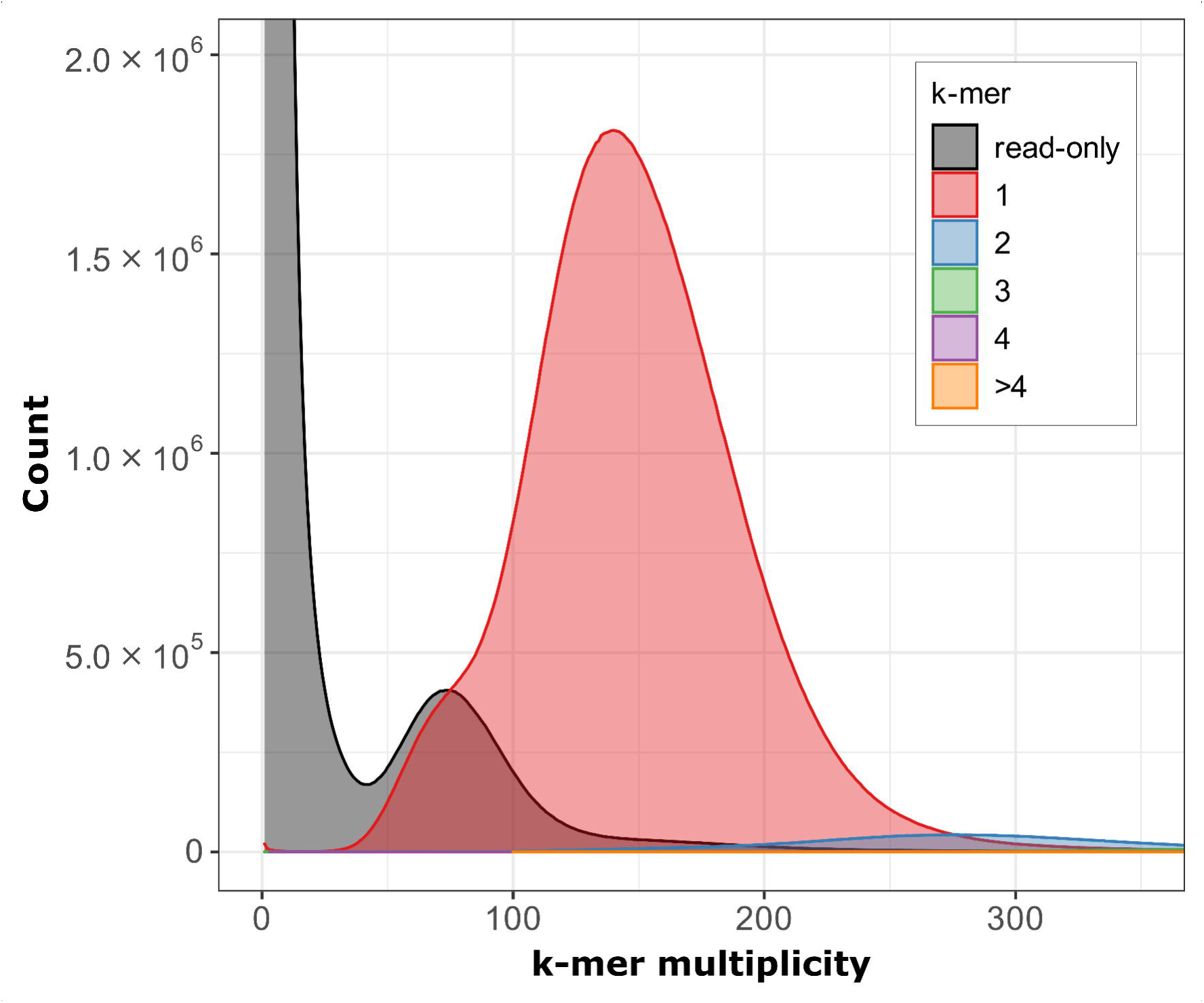
Genome assembly of *Adelges tsugae*, ihAdeTsug: Merqury assembly spectrum plot. Multiplicity of each *k*-mer found in the 10x Genomics Chromium linked-read set, coloured by the number of times it is found in the *A. tsugae* (ihAdeTsug) assembly.

Comparisons of chromosome synteny between *A. tsugae* (ihAdeTsug), *S. chinensis*, and *R. maidis* using ChromSyn revealed strong conservation of sex chromosome composition and structure across the Aphidomorpha (**Figure 6**), as has been previously demonstrated without comparisons including a chromosome-scale adelgid assembly^60, 63, 64, 65^. Two scaffolds were syntenic with sex chromosome(s) previously identified in the genomes of the two other taxa^60^, providing confirmation of a 2(X_1_X_2_)/X_1_X_2_0 (♀/♂) sex determination system for *A. tsugae.* These are identified in the *A. tsugae* (ihAdeTsug) assembly as chromosomes X1 and X2. The presence of multiple sex chromosomes within adelgid species has been demonstrated previously^22, 64^. Similarly, there was strong concordance between the eight remaining autosomes for *A. tsugae* and the ten identified in *S. chinensis,* suggesting that elevated autosomal rearrangement within the Aphididae^64^ may be limited to aphid subfamilies that have undergone more recent diversification.

**Figure 6.**
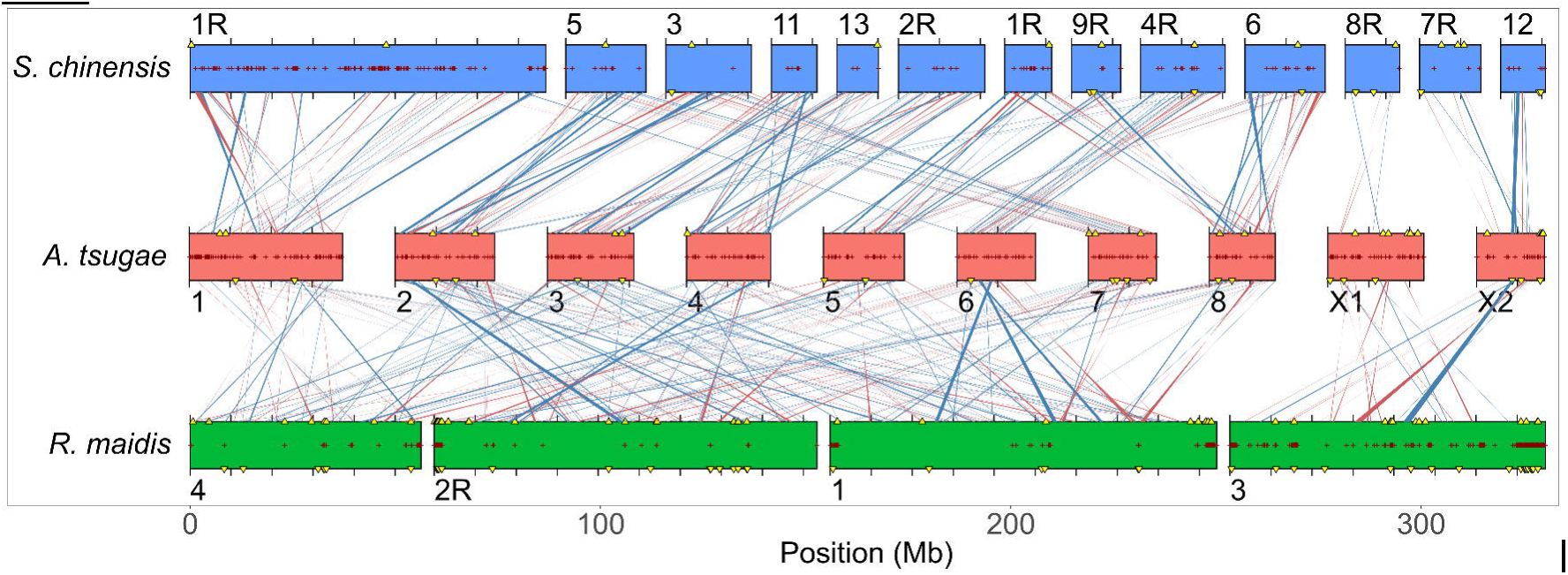
Comparison of chromosome synteny between *Adelges tsugae* (ihAdeTsug), *Schlechtendalia chinensis*, and *Rhopalosiphum maidis*. Three-way ChromSyn plot showing connections between syntenic regions greater than 50 Kb between the assemblies. Red lines are inverted regions. Plus symbols within chromosomes denote assembly gaps, and ticks along the chromosomes are spaced every 10 Mb. Yellow triangles are predicted positions of BUSCO orthologs that are not found in the *A. tsugae* assembly, based on assumptions of conserved synteny with *S. chinensis* and *R. maidis*. Location of triangles above or below the chromosome centre-line indicates the strand. Reversed chromosomes are labelled with an “R” suffix. Chromosomes 7, 8, and 12 and chromosome 3 are the sex chromosomes of *S. chinensis* and *R. maidis*, respectively.

The mitochondrial genome was also assembled into a completely circular contig 25,980 bp in length (**Figure 7**). It appears as the last entry in the multi-fasta file of the NCBI genome submission (GCA_045999825.2). Despite the discrepancy in length from the existing HWA mitogenome reference (MT263947)^53^, the length reported here is consistent with at least two other recently published adelgid and aphid mitogenomes^22,66^. The arrangement of protein-coding genes (PCGs), transfer RNAs (tRNAs), and ribosomal RNAs (rRNAs) is also similar between the two HWA mitogenomes, with approximately 94% pairwise sequence identity among all PCGs and both rRNAs, as expected based on prior characterizations of this species’ global genetic diversity^3^. The mitogenome consists of 13 PCGs, 22 tRNAs, two rRNAs, and large control (4655 bp) and repeat (5755 bp) regions characterized by AT-rich tandem repeats. The control region contains an imperfect compound repeat of 232 bases (2.9 copies) plus 23 bases (2 copies) with a 3 bp overlap repeated four times between tRNA-Ile and rRNA-S. The repeat region is composed of an imperfect repeat of 207 bases with 24.9 copies between nucleotide positions 420 and 6175.

**Figure 7.**
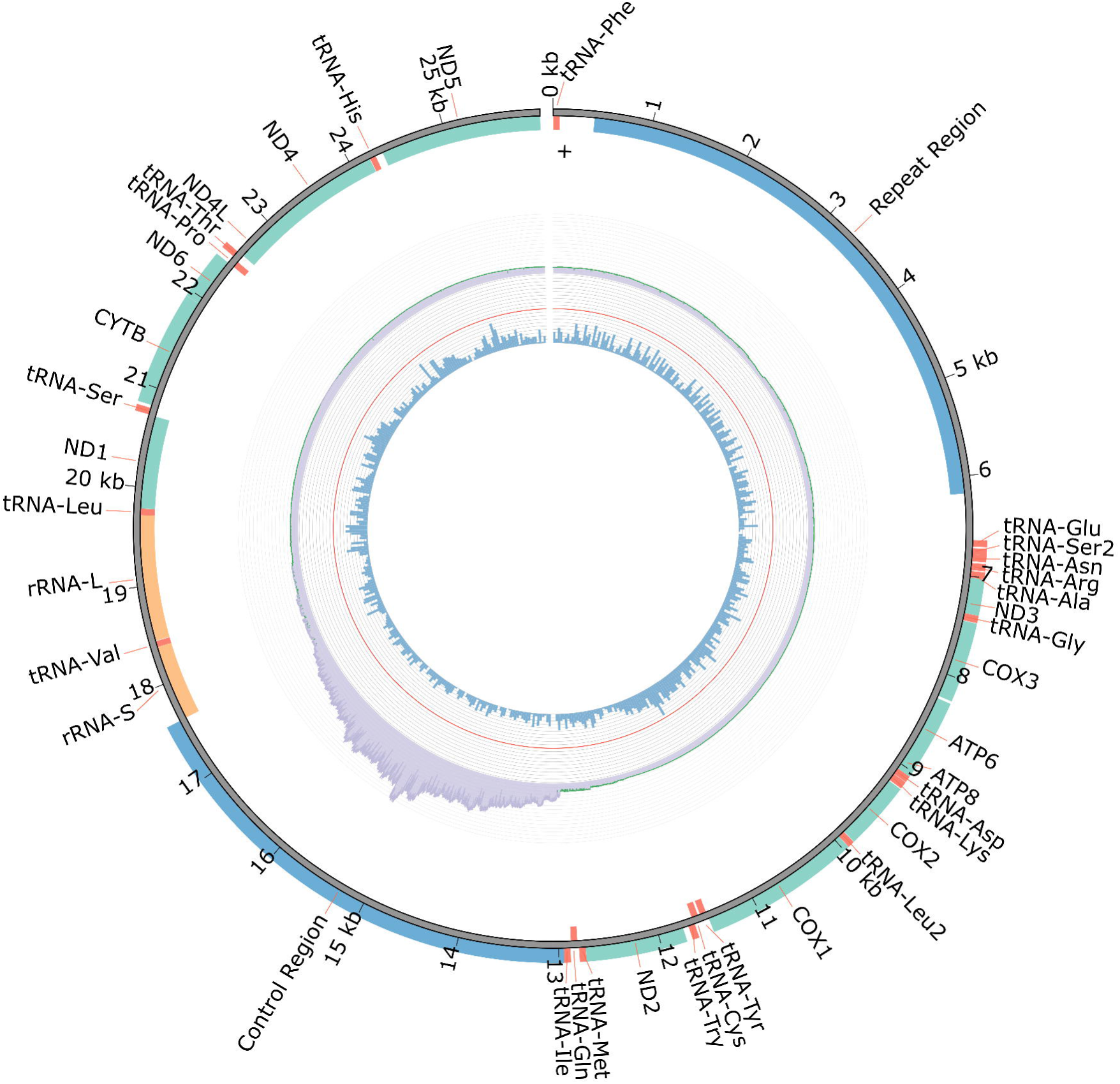
Mitochondrial genome assembly of *Adelges tsugae*, ihAdeTsug. Circos plot of *A. tsugae* (ihAdeTsug) mitochondrial genome assembly, generated from the visualize command of MitoZ. The location of protein-coding genes (green-cyan), tRNAs (red), rRNAs (orange), and the control and repeat regions (blue) are displayed on the outer track, with orientation denoted by placement above or below. Read depth of pre-assembly reads from Falcon used in the mitogenome assembly is displayed on the middle track. GC composition is displayed on the inner track using a window size of 50 bp.

### Genome Annotation

Genome annotation was performed by the European Bioinformatics Institute using the Ensembl Gene Annotation system^67^ (https://beta.ensembl.org/help/articles/non-vertebrate-genome-annotation) and publicly available RNA-seq data on NCBI (PRJNA242203). The *A. tsugae* (ihAdeTsug) assembly is composed of 11,800 coding genes, 1930 non-coding genes and 20,403 mRNA transcripts.

## Data availability

Genome data is available from NCBI’s Sequence Read Archive (SRA): *Adelges tsugae*; accession numbers SRX27535639 (long-reads), SRX27535640 (short-reads), SRX27535641 (linked-reads), and SRX27535642 (Hi-C); and the assembled genome is available from NCBI’s Assembly database: *Adelges tsugae*; accession number GCA_045999825.2. All data are linked under BioProject PRJNA1187642.

## Author contributions

Bryan Brunet – conceptualization, data curation, formal analysis, funding acquisition, investigation, methodology, project administration, visualization, writing – original draft preparation, writing – review & editing

Dustin Dial – formal analysis, data curation, methodology, writing – review & editing

Gaelen Burke – funding acquisition, investigation, resources, writing – review & editing

Carol von Dohlen – funding acquisition, investigation, resources, writing – review & editing

Julia Douglas Freitas – formal analysis, methodology, writing – review & editing

Haley Sanderson – formal analysis, methodology, writing – review & editing

Afiya Chida – data curation, formal analysis, writing – review & editing

Samantha Jones – data curation, writing – review & editing

Fergal Martin – data curation, writing – review & editing

Leanne Haggerty – data curation, writing – review & editing

Stephen Scherer – conceptualization, funding acquisition, writing – review & editing

Ioannis Ragoussis – conceptualization, funding acquisition

Steven Jones – conceptualization, funding acquisition, writing – review & editing

Robert Foottit – conceptualization, writing – review & editing

Nathan Havill – conceptualization, funding acquisition, investigation, resources, writing – review & editing

## Acknowledgements

The authors would like to acknowledge several other people for their contributions to this work: Michael Montgomery collected sample# 19-124-03 which was used in the long-read sequencing libraries; Jackson Eyres provided assistance in coordinating bioinformatic expertise at Agriculture and Agri-Food Canada’s Ottawa Research and Development Centre; and J. Spencer Johnston performed the genome size estimates by flow cytometry at the Department of Entomology, Texas A&M University, College Station, Texas, USA. The authors would also like to acknowledge the University of Toronto McLaughlin Centre, and S. Scherer’s endowed position as Northbridge Chair in Paediatric Research at The Hospital for Sick Children (Toronto, Canada).

